# Importance of Apparatus Scaling in Novel Object Recognition for Juvenile and Adult Rats

**DOI:** 10.1101/2025.11.17.683970

**Authors:** Laura Baron, Samantha Hetherington, Andrew M. Poulos

**Affiliations:** Department of Psychology and Center for Neuroscience Research, University at Albany, Albany, New York 12222

**Author notes:** **Corresponding Author:** Dr. Andrew M. Poulos, Department of Psychology, University at Albany, State University of New York, 1400, Washington Avenue, Albany, New York, 12222.

## Abstract

The growing interest in ontogenetic studies of learning and memory, along with early-life perturbations, has led to the use of younger rodents as a key biological variable in many investigations. This development prompts an important question about whether procedures and apparatuses designed for studying learning and memory in adults should be simply adapted for use with younger and smaller rodents. The current study examined how arena size affects novel object recognition (NOR) performance in juvenile and adult rats. A commonly used larger arena reliably detected novel object preference in adults but not in juveniles. Adjusting the arena size based on average weight differences between age groups improved the consistency of NOR performance in juvenile rats. Sex differences were observed: adult males performed reliably across all arena sizes, whereas adult females required larger arenas to demonstrate effective NOR performance. These findings highlight the importance of tailoring arena dimensions to developmental stage and sex for accurate cognitive assessment. Specifically, they support the use of weight-based arena scaling as a methodological approach in developmental neurobehavioral research and emphasize the importance of careful design when studying female rodents. Future studies should explore similar environmental adjustments for other behavioral tests in juvenile and female populations.

## Introduction

Most behavioral assays for learning have been established and validated to assess learning in adult male rodents (Ennaceur & Delacour, 1988), which may present unique challenges in studying learning and memory in developing animals. A widely used learning paradigm, the Novel Object Recognition (NOR) task, originally developed in adult male Wistar rats to assess recognition memory within an environmental apparatus, relies on an animal’s innate preference for novel objects over familiar ones (Ennaceur & Delacour, 1988). It remains widely used in primates (Neiworth et al., 2023), mice (Gao et al., 2025), and rats (García-Aviles et al., 2025) to explore the cognitive and neural mechanisms of learning and memory.

Recognition memory, as evaluated by NOR, relies on medial temporal lobe regions of the brain that undergo an extensive maturation process concurrent with significant physical growth. This underscores a vital consideration in the study of the ontogenetic development of learning: environmental factors, such as the dimensions of the training apparatus and object stimuli, may differentially influence performance across developmental stages. Consequently, a critical aspect in assessing learning in juvenile subjects is whether the apparatuses commonly calibrated for adults are suitable for smaller, younger individuals.

In recent years, standard NOR procedures typically involve habituation to an empty open-field arena, exposure to identical objects during familiarization training, and subsequent testing with one familiar and one novel object following a retention interval (Lueptow, 2017). Exploration time directed toward the novel versus familiar object is used as an index of recognition memory. Although these procedures are generally consistent across laboratories, the size of both the arena and objects varies considerably (Table 1). For adult rodents, such variation may not be particularly consequential; however, for juveniles, large open arenas may diminish exploratory and object investigatory behavior due to competing anxiety-like behaviors. Moreover, object size may constrain encoding if smaller animals cannot adequately investigate object stimuli. Both factors may therefore confound the performance in developmental studies of novel object recognition.

Despite the widespread application of NOR, limited research has systematically investigated whether scaling the apparatus dimensions to the body size of juvenile animals affects learning outcomes. Some studies have employed adult-sized arenas for juveniles (Jablonski et al., 2013; Ramsaran et al., 2016; Westbrook et al., 2014), while others have adopted smaller, mouse-sized apparatuses (Bates & Trujillo, 2023; Socha et al., 2024; Witek et al., 2023). The latter approach, however, may reflect convenience rather than measured scaling based on body mass and developmental stage (Reger et al., 2009). Although studies have constructed experimental settings with this consideration in mind (Jablonski et al., 2013; Reger et al., 2009), such variations may hinder cross-study comparisons and constrain the interpretability of findings regarding the development of recognition memory.

The current study addresses this gap by evaluating the influence of arena and object size on NOR performance in juvenile and adult male and female Long-Evans rats. We constructed two arenas scaled to the average body mass of postnatal day (P) 24 (79 g) and 90 (300 g) rats: a large, standard 1.0 m^2^ arena (LNOR) and a smaller 0.26 m^2^ arena (SNOR). Objects were similarly scaled in height and volume in correspondence to arena sizes. We hypothesized that juvenile rats would show greater NOR in the SNOR compared to the LNOR, reflecting improved object engagement. In contrast, we predicted that adults would exhibit NOR regardless of apparatus size. By experimentally manipulating the apparatus dimension, this study aims to clarify methodological considerations for developmental research using NOR and to improve the reliability of recognition memory measures in rodents across the lifespan.

## Methods

### Subjects

Male and female Long-Evans rats were bred in-house at the University of Albany, State University of New York, animal facility, and were housed in groups of 2-4 in same-sex clear cages. Pups were weaned on postnatal day 21. The colony room was temperature- and humidity-controlled and maintained on a 14:10 hour light-dark cycle, with all experiments conducted during the light phase. All animals had access to food (Prolab® IsoPro® RMH 3000, ScottPharma, Inc., Marlborough, MA) and water ad libitum. All animal care and experiments were carried out on the University at Albany campus in accordance with the Institutional Animal Care and Use Committee (IACUC) at the University at Albany, SUNY. Total number of subjects per group: Juvenile Large Arena, Male n=14, Female n=11; Juvenile Small Arena, Male n=11, Female n=16; Adult Large Arena, Male n=23, Female n=13; Small Arena, Male n=14, Female n=17.

### Handling and habituation

All animals were handled for 3-4 minutes daily for three consecutive days prior to behavioral testing. On testing days, animals were acclimated to the holding room for 15-30 minutes before behavioral testing. 24 hours before object exposure, animals were placed into a clean and dry arena for 15 minutes (**Figure 2, left panel**). Two arenas were constructed using PVC board, adhered with thermoplastic adhesive, and laminated in blue contact paper. LNOR, the large arena, measured at 100cm x 100cm x 40cm (LxWxH). SNOR, the small arena, measured at 51cm x 51cm x 30cm. (see **Figure 1**), and animal placement was determined by randomized group assignment. Luminance at the center of the arenas was 21 lux for the SNOR and 24 lx for the LNOR, with all four corners of both arenas at 11 lux. This luminance at the edges was the darkest to obtain video files with optimal visual accuracy. A camera (SONY HDR-CX405, Tokyo, Japan) was positioned overhead via a tripod secured to the ceiling directly above the testing arena. Arenas were cleaned with 70% ethanol between sessions and disinfected with Clidox at the end of each day of testing.

**Figure 1.**
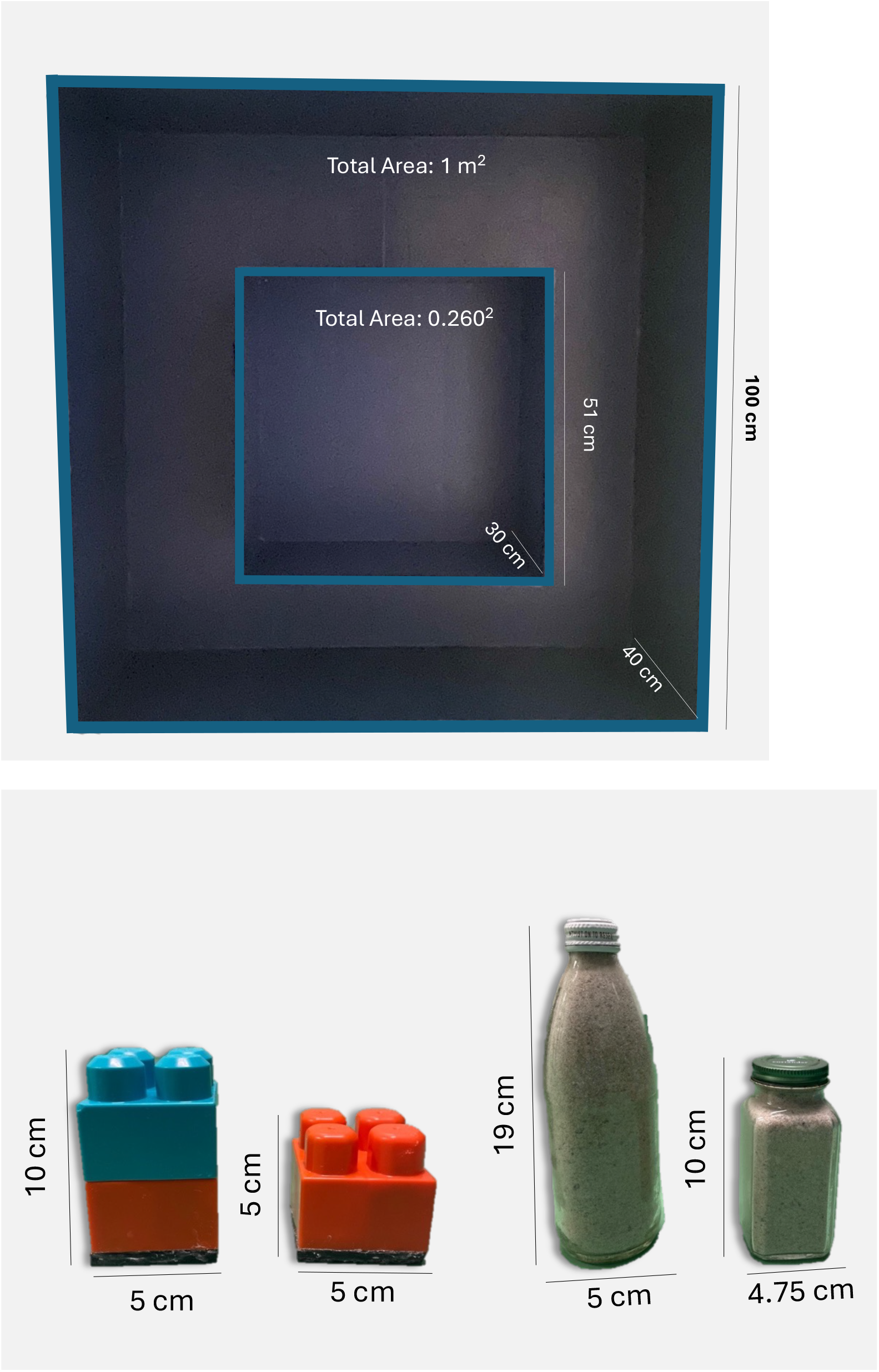

**Figure 2.**
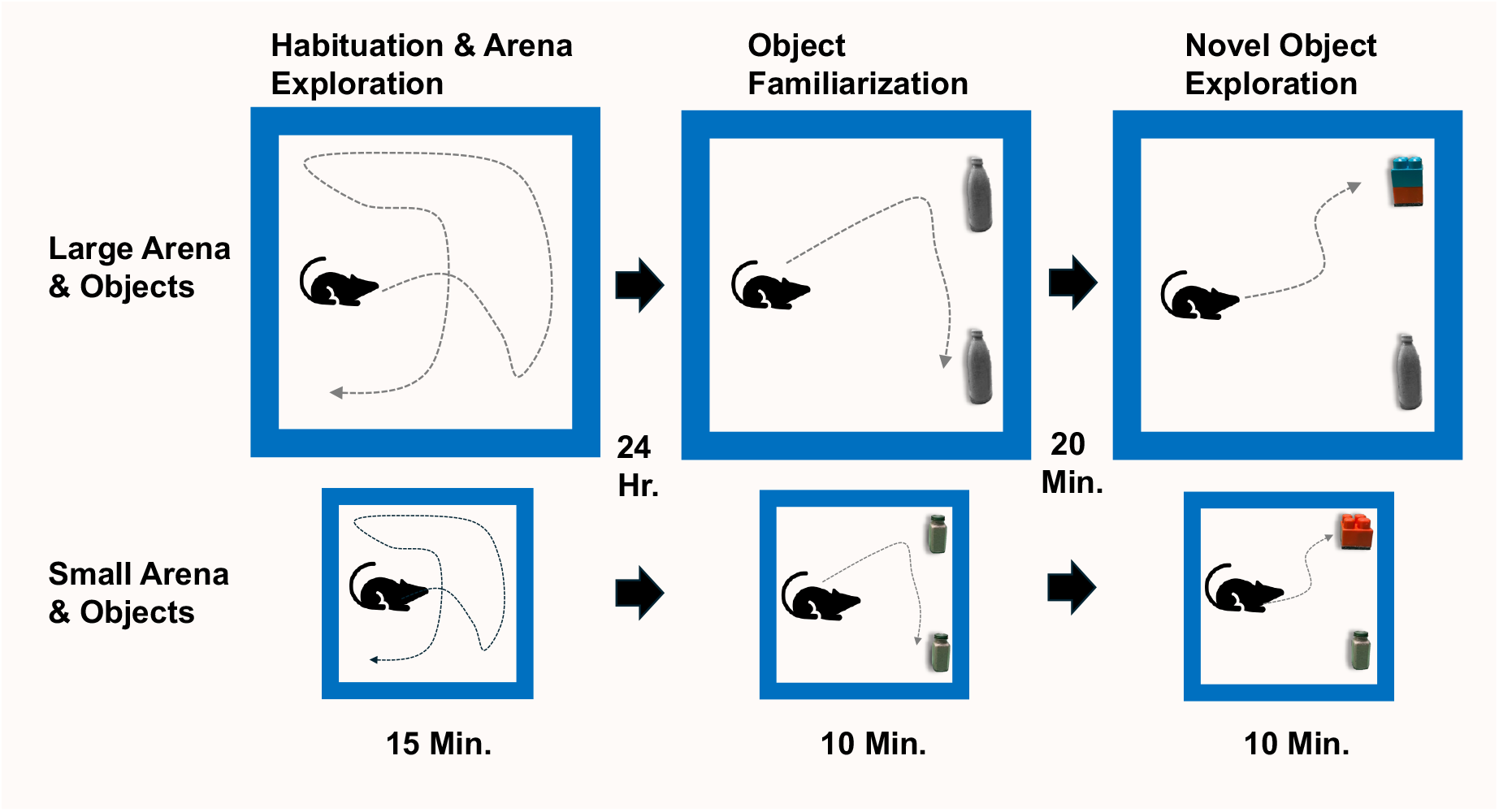

**Figure 3.**
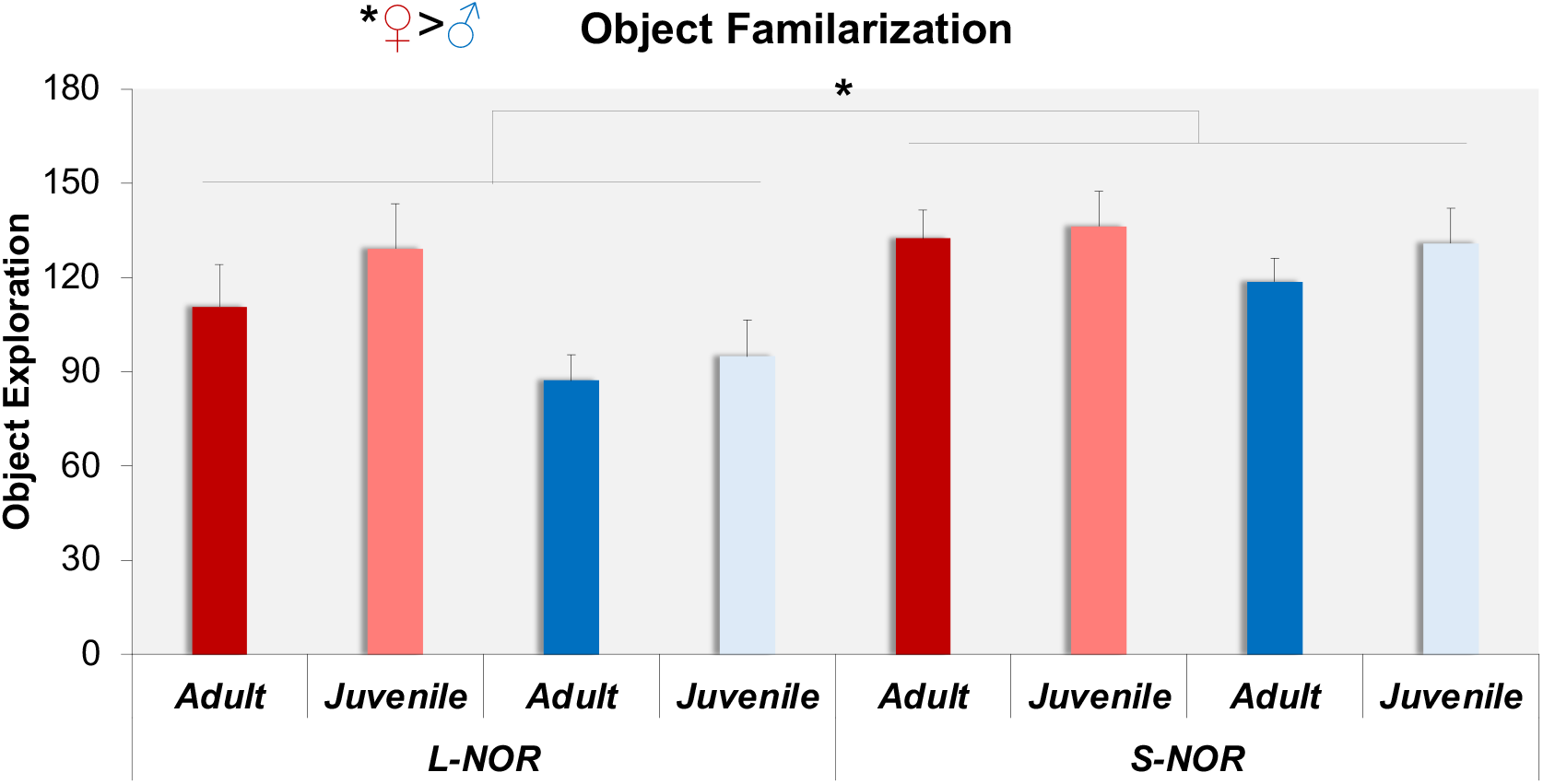

### Familiarization

24 hours after pre-exposure to the arena, animals were returned to the arena and presented with two novel objects in each corner of the wall opposite the rats’ starting point, 5 cm away from each wall (**Figure 2**, middle panel). For the SNOR, the objects were either two 4-oz glass jars or two single-platform Mega Bloks (Mattel, El Segundo, CA) (10.16 x 10.16 x 5.08 cm). For the LNOR, the objects were either two 12-oz glass bottles or a double-stack platform of Mega Bloks (10.16 x 10.16 x 10.16 cm) (see Figure 1). All objects were filled with sand for weight and sealed. Animals were allowed to explore the arena and the objects for 10 minutes.

### Recognition Test

After a twenty-minute period, short-term memory was evaluated by returning the animal to the arena, where one of the two familiar objects was replaced with a novel object (**Figure 2**, right panel). The sequence and presentation of objects (i.e., whether the novel object was positioned on the left or the right) were counterbalanced across trials. The animals were permitted to explore the arena and the objects for a duration of ten minutes. The arenas were cleaned with 70% ethanol between sessions and disinfected with Clidox after each testing day. *Behavioral Scoring:* Increased interaction with the novel object compared to the familiar object (greater than 50% of total investigation time) indicated successful short-term memory encoding. Object investigation was scored by an observer blind to the experimental design and was quantified as active nose-pointing within three centimeters of the object’s perimeter.

### Statistical Analysis

To determine whether object learning occurred within each group, individual group means were compared to 50% using a single-sample Student’s T-Test. To determine between-group differences, an overall 3-way ANOVA (age x arena size x sex) was performed (α=.05).

## Results

### Object Familiarization

During the familiarization period, animals spent significantly more time exploring objects in the SNOR compared to the large standard environment **(Figure 4)**. Notably, females spent more time exploring objects overall, but this did not affect encoding, as shown during the recognition test **(Figure 5)**. An ANOVA revealed a significant main effects of environmental size *F* (1,90) = 4.706, *p*<.05), where animals in the SNOR (M = 130 seconds, SD = 36.3), spent more time exploring objects stimuli than animals in LNOR (M = 102 seconds, SD = 45.5), and sex *F* (1,90) = 6.789, *p*<.05) where females (M = 128 seconds, SD = 43.5) explored more than males (M = 104 seconds, SD = 40.5), but no effect of age *F* (1,90) = .616, *p* >.05). There were no interactions between sex and arena size F (1,90) =.395, p >.05), sex and age (F (1,90) =1.859, p >.05), size and age F (1,90) =.548, p >.05), or among sex, size, and age F (1,90) =.120, p >.05).

**Figure 4.**
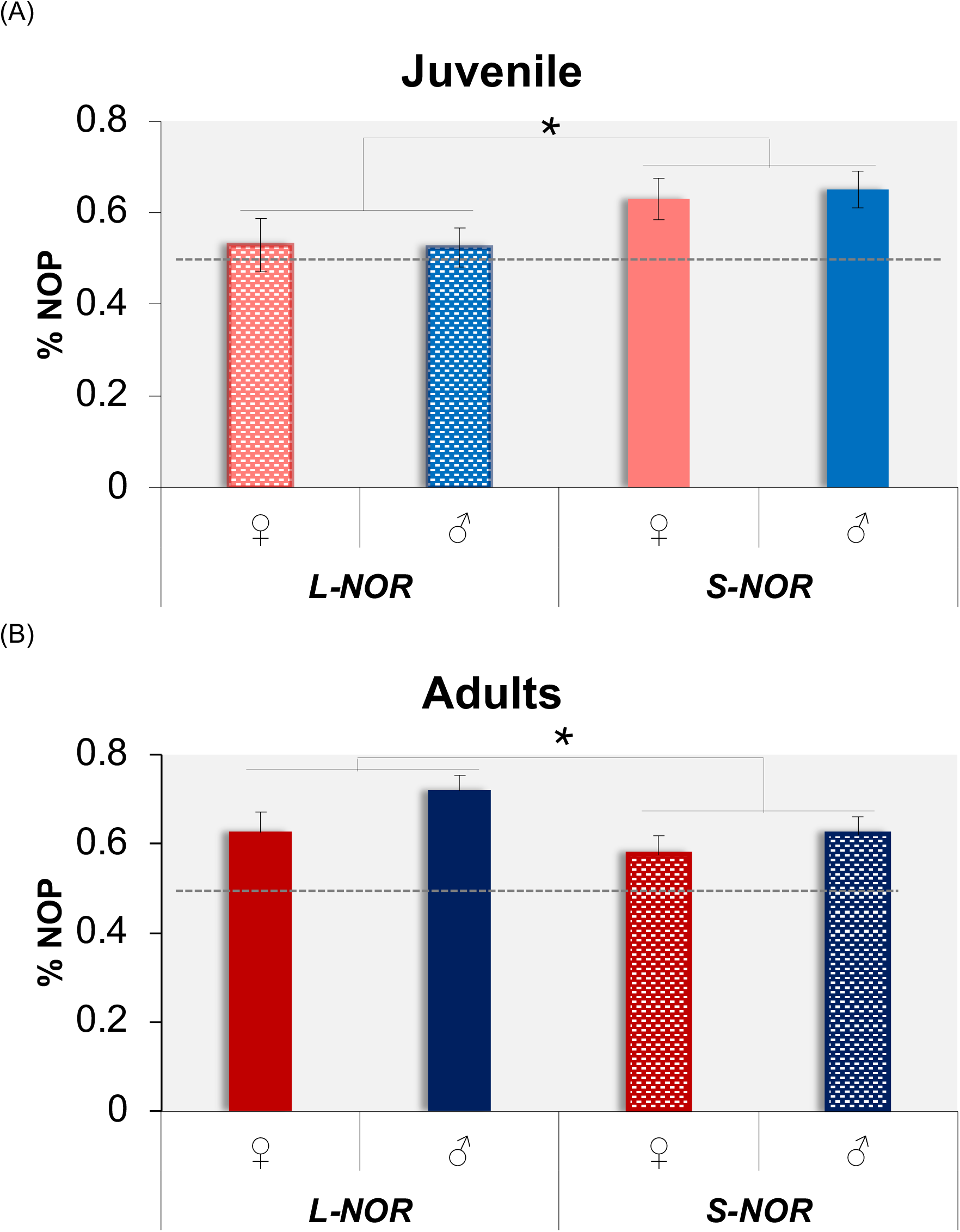

**Figure 5.**
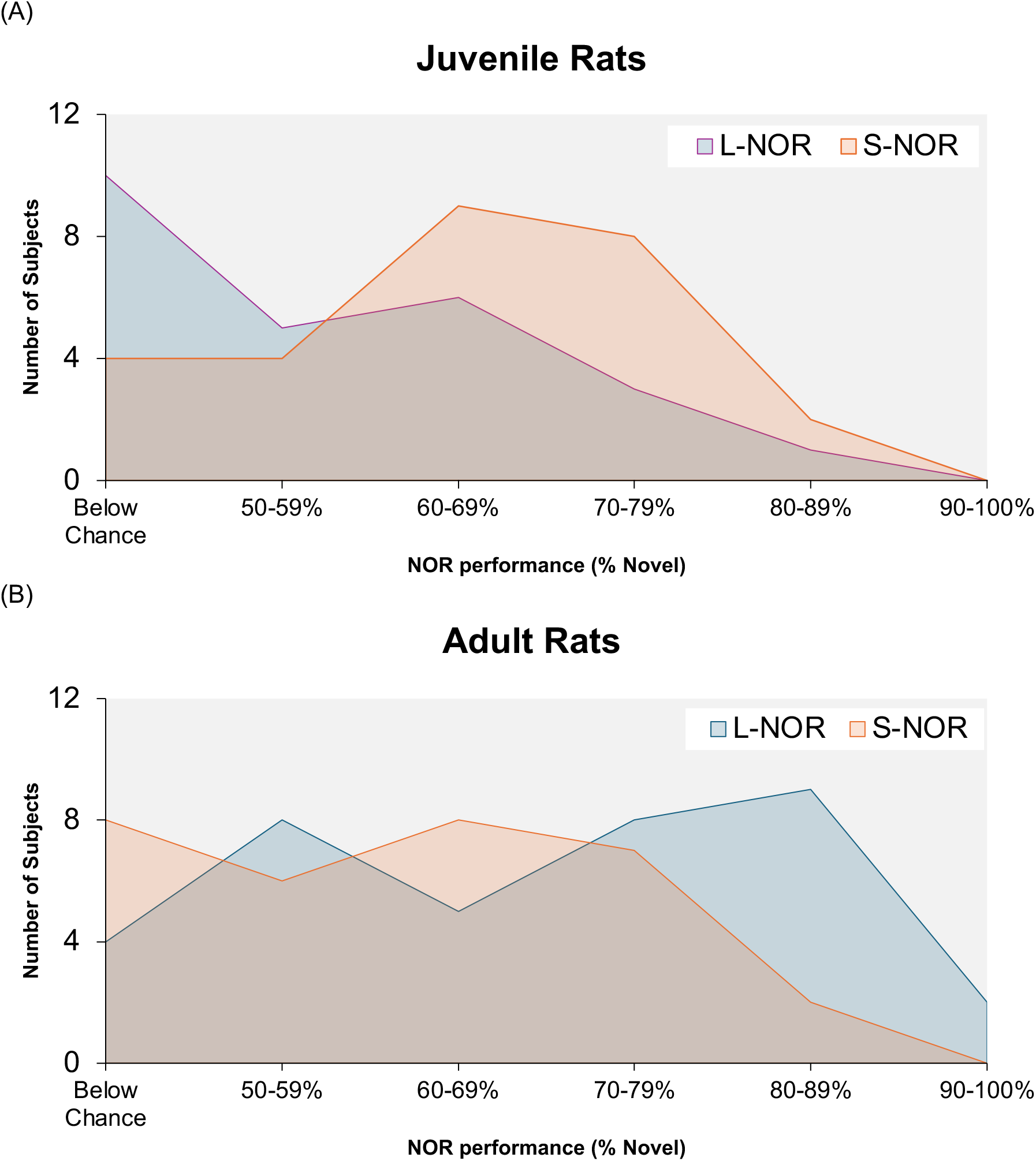

### Novel Object Recognition Test

When the arena size was adjusted relative to juvenile or adult body weight, specifically for the large-to-adult and small-to-juvenile conditions, it produced consistent and reliable preferences for novel objects in both juvenile and adult rats. However, when the arena size was not aligned with body weight, specifically for small-to-adult and large-to-juvenile comparisons, novel object preference was not evident in juvenile males and females (**Figure 5A**), as well as in adult females, but was not reliable in adult males (**Figure 5B**). A one-sample T-test compared to chance levels (50%) revealed that age-matched arena sizes are generally necessary for supporting object learning. Specifically, adult females showed a NOP in the LNOR (M = 62.7%, SD = .162, t(12)=2.819, p<.01) but not in the SNOR (M = 57.7%, SD = .169, t(17)=.887, p >.05), while adult males exhibited NOP in both the LNOR (M = 72%, SD = .161, t(23)=6.754, p<.001) and the SNOR (M = 62.1%, SD= .153, t(13)=2.965, p<.01). In juvenile rats, both females and males showed a NOP in the SNOR (female: M = 63%, SD = .184, t(16) = 3.178, p < .01; male: M = 65.1%, SD = .132, t(10) = 3.791, p < .01) but not in the LNOR (female: M = 52.8%, SD = .191, t(10) = .493, p > .05; male: M = 52.3%, SD = .159, t(12) = .679, p > .05) (**Figure 5**).

To test for contributions of arena and object scaling in juvenile and adult male and female rats, a three-way analysis of variance showed no significant main effects of age, sex, or arena size. However, it revealed a significant interaction between age and arena size, F (1, 120) = 9.992, p < .01, indicating that NOP was higher in the larger arena for adults, whereas it was higher in the smaller arena for juveniles. Post hoc analyses with Bonferroni-corrected T-tests confirmed this: in the LNOR arena, adult rats (M = 68.6%, SD = .166) had significantly greater NOP than juveniles (M = 52.6%, SD = .17, p < .001) (**Figure 5**), and juvenile animals showed significantly higher NOP in the small arena (M = 63.9%, SD = .162) compared to the large arena (M = 52.6%, SD = .17, p < .05). These results suggest that NOR performance is optimal when juvenile and adult rats are tested in an arena environment with objects proportionate to their developmental size.

The contribution of apparatus size to juvenile and adult NOP performance is evident in Figure 6, where animal performance is distributed across 10% bins of novel object preference. Juvenile rats in SNOR exhibited a non-uniform inverted U-shaped distribution, with most scores peaking around 60 to 79% with 19 of 27 subjects (70.4%) exhibiting NOP above 60%, while only 4 subjects (14.8%) showed performance below 50% chance levels (**Figure 6A**). Conversely, in the LNOR, juvenile rat NOP performance was skewed to the right, with 10 of 25 (40%) juvenile animals not demonstrating learning (NOP below 50%), and just 10 juvenile subjects showed learning above 60%. In contrast to juveniles, adult rats in LNOR showed a bimodal distribution with peak NOP scores around 70 to 89% and 50-59% with 4 of 36 (11.1%) subjects failing to demonstrate learning, while 24 subjects (66.7%) showed NOP above 60% (**Figure 6B**). In SNOR, the relative distribution of scores is skewed to the right, as 8 of 31 adult rats (25.8%) did not show NOP, and 17 subjects (54.8%) exhibited learning above 60% NOP, yet only two rats scored above 80%. These data indicate that higher NOR scores predominate when juvenile animals are tested in an environment scaled to their relative size, in a smaller rather than a larger NOR apparatus.

Lastly, a Pearson correlation coefficient was used to assess the linear relationship between the time spent investigating objects during the familiarization phase and the recognition phase, which was not significantly correlated (r (90) = -0.078, p > 0.05).

## Discussion

The present study examined the effects of arena and object scaling on novel object recognition (NOR) in juvenile and adult Long-Evans Hooded rats. Our findings demonstrate that apparatus dimensions relative to animal weight critically influence recognition memory performance. Specifically, juvenile rats exhibited robust novel object preferences (NOP) only in the scaled-down environment. In contrast, adults showed consistent NOP in the standard large arena, with more variable outcomes in the scaled-down apparatus. These results suggest that the physical scale of the testing environment, in relation to the developmental stage of the animal, plays a vital role in supporting object encoding and recognition.

A primary finding of this study is that juvenile animals failed to reliably exhibit NOP when tested in the larger arena commonly used for adult rodents. Instead, they showed enhanced exploration and greater NOP in the scaled apparatus, in which the arena and object sizes were matched to body weight. This aligns with previous reports highlighting the importance of considering developmental constraints in designing behavioral assays (Reger et al., 2009; Bates & Trujillo, 2023). Larger open environments are well known to elicit thigmotaxis and risk-assessment behaviors (Denenberg, 1969; Palm et al., 2014), which may compete or suppress exploratory behaviors required for object encoding (Jablonski et al., 2013). To minimize this in the current study, an extended 20-minute habituation preceded the familiarization session. Our findings that juveniles spent more time investigating objects in the scaled arena further support the interpretation that a proportionally scaled environment facilitates engagement with object stimuli.

In contrast, adult animals performed reliably in the more standard large arena, consistent with the task’s original design and validation in adult rodents (Ennaceur & Delacour, 1988; Reger et al., 2009). Interestingly, adults also displayed NOP in the smaller scaled apparatus, though performance distributions suggested weaker and more variable recognition compared to the larger arena. This may reflect the fact that adults are sufficiently motivated and capable of engaging with objects across a range of contextual environments, though suboptimal scaling may reduce the robustness of encoding. Collectively, these findings indicate that apparatus scaling is more critical for juveniles than for adults, with developmental stage determining the optimal match between rodent size and testing environments.

Sex differences also emerged in overall object exploration during familiarization, with females investigating more than males, but this effect did not translate into superior recognition performance. This aligns with prior research showing that while sex can influence exploratory behavior and anxiety-like responses (Levy et al., 2023; Shansky, 2018; Witek et al., 2023), it does not consistently alter NOR performance (Becegato & Silva, 2022; but see Cost et al., 2012). Notably, both male and female juveniles required scaled environments for reliable learning, highlighting that developmental stage rather than sex is the key determinant of task performance under the current experimental conditions.

These findings have important methodological implications for NOR procedures. First, they suggest that a one-size-fits-all approach to NOR testing across development may confound interpretation of learning and memory. The failure of juveniles to demonstrate NOR in adult-sized arenas does not necessarily indicate immature recognition learning or memory but may instead reflect inappropriate scaling. Second, scaling the arena and its objects may prove to be crucial, as objects that are disproportionately large relative to the animal’s body size could have limited tactile and visual exploration, thereby impairing encoding. Thus, consideration of apparatus scaling is crucial for developmental studies of recognition memory and for improving cross-study comparability.

There are some limitations of the current study that should be acknowledged. Although we tested both sexes and two developmental periods, additional ages would provide a more detailed trajectory of how scaling requirement changes more broadly across development (Jablonski et al., 2013; Westbrook et al., 2014). In addition, while anxiety-like behavior is a potential confound that could influence exploratory patterns, we did not directly measure risk assessment or thigmotaxis; inclusion of these measures could strengthen future interpretations of how stress and affect interact with learning in differently sized arenas. Finally, while the current study focused on scaling arena and object dimensions by body weight, future work should consider whether other factors, such as sensory-motor maturation (Rudy & Paylor, 1987; Schenk & Morris, 1985) and stress-reactivity, interact with environmental design to influence NOR. Indeed, it has been demonstrated that juvenile rats exhibit an enhanced hypothalamic-pituitary-adrenal axis response to stress compared to adults (Klein & Romeo, 2013).

In conclusion, our results show that novel object recognition performance in rats is strongly influenced by the proportional relationship between animal size and testing apparatus. Juvenile rats require scaled-down environments to exhibit reliable learning, whereas adults perform robustly in standard arenas but show reduced performance in smaller environments. These findings emphasize the necessity of adjusting experimental parameters when investigating learning and memory across development and suggest that scaling behavioral apparatuses to animal sizes is an essential factor in achieving valid and interpretable outcomes.

